# Reporting amyloid beta levels *via* bioluminescence imaging with amyloid reservoirs in Alzheimer’s disease models

**DOI:** 10.1101/2021.06.05.447217

**Authors:** Jing Yang, Weihua Ding, Biyue Zhu, Sherri Zhen, Shi Kuang, Can Zhang, Peng Wang, Fan Yang, Liuyue Yang, Wei Yin, Rudolph E. Tanzi, Shiqian Shen, Chongzhao Ran

## Abstract

Bioluminescence imaging has changed daily practice in preclinical research of cancers and other diseases in the last decades; however, it has been rarely applied in preclinical research of Alzheimer’s disease (AD). In this report, we demonstrated that bioluminescence imaging could be used to report the levels of amyloid beta (Aβ) species in vivo. We hypothesized that AkaLumine, a newly discovered substrate for luciferase, could bind to Aβ aggregates and plaques. We further speculated that the Aβ species have the reservoir capacity to sequester and release AkaLumine to control the bioluminescence intensity, which could be used to report the levels of Aβs. Our hypotheses have been validated *via in vitro* solution tests, mimic studies with brain tissues and mice, two-photon imaging with AD mice, and *in vivo* bioluminescence imaging using transgenic AD mice that were virally transduced with aka Luciferase (AkaLuc), a new luciferase that generates bioluminescence in the near infrared window. As expected, compared to the control group, we observed that the Aβ group showed lower bioluminescence intensity due to AkaLumine sequestering at early time points, while higher intensity due to AkaLumine releasing at later time points. Lastly, we demonstrated that this method could be used to monitor AD progression and therapeutic effectiveness of avagacestat, a well-studied gamma-secretase inhibitor. Importantly, a good correlation (R^2^ = 0.81) was established between in vivo bioluminescence signals and Aβ burdens of the tested AD mice. We believe that our approach can be easily implemented into daily imaging experiments and has tremendous potential to change daily practice of preclinical AD research.

## Introduction

No cure and effective drug treatment are available for Alzheimer’s disease (AD) so far ^1-5^. Clearly, drug development for AD is an urgent task. Compared to other diseases such as cancers, however, AD drug development both at the clinical and preclinical stages is far behind ^6,7^. The number of AD therapeutics under preclinical discovery is unquestionably dwarfed by the number of cancer drugs at the similar stages ^7,8^. Among numerous factors that have contributed to the stagnant state of AD drug development, the shortage of simple but effective methods for monitoring therapeutic effects has significantly impeded the progress of preclinical studies.

There is nearly no argument that tumor mass (volume) is one of the most important and direct parameters to assess the effectiveness of drug treatment in preclinical research, clinical studies, and patient treatment. Shrinking and reducing tumor mass represent the greatest challenge, regardless of the pathways of drug targeting. In fact, nearly all anti-cancer drugs are aimed to reduce the tumor mass ^9,10^. With a parallel comparison, we may consider that the levels of Aβ and tau in AD research are “equal” to tumor mass in cancer research. Similarly, regardless of drug targeting pathways, reducing Aβ and Tau levels is one of the most essential aims (may not be the sole aim) for AD drug development. However, only a few tools are available to assess the level of Aβ and tau levels in vivo, compared with plentiful tool arsenal for reporting tumor mass. Bioluminescence imaging (BLI) has been used in thousands of biomedical laboratories for preclinical cancer research, due to its high robustness, excellent signal-to-noise ratio and easy-to-use. By contrast, such a powerful imaging technology has rarely been used in preclinical research of Alzheimer’s disease ^11^.

Behavioral testing has been often used to evaluate the therapeutic effects of AD drug candidates ^12,13^; however, this approach is not only tedious and time consuming, but also can’t accurately reflect the amounts of amyloid beta (Aβ) deposits and tau tangles ^14,15^, the most relevant molecular biomarkers for AD pathology ^16^. Although positron emission tomography (PET) and magnet resonance (MR) imaging have the capacity to monitor AD pathology in human studies and animal studies, expensive equipment, high cost and the need of well-trained experts are their limitations ^17,18^. By contrast, BLI is easy-to-use and low cost. More importantly, it provides high signal-to-noise ratio (SNR) ^19^, which is crucial for reliable quantification. For tumor imaging, luciferases, the enzyme for generating bioluminescence, can be easily engineered into the target of interest without interrupting the properties and functions of the target ^20-22^. However, protein engineering of Aβ via inserting a luciferase is essentially impracticable, because this will drastically change their properties and functions. Aβ is not compatible for tagging a 60KD luciferase ^23^, due to its much smaller molecular weight (4KD). Moreover, the tagging of luciferase can profoundly change Aβ’s physiological behaviors such as aggregating and trafficking. Tau proteins are larger sizes and have been co-expressed with luciferase ^11,24^. However, inserting luciferase into a tau protein could potentially alter its functions and aggregation dynamics as well. Clearly, these factors have been the major hurdles to utilize BLI for preclinical AD research. Several groups attempted to establish BLI methods for Aβ reporting via co-expressing luciferase with amyloid precursor protein (APP) in AD mouse models ^25-28^; however, these methods have been rarely used to accurately report the levels of Aβs, because the level of APP is not equal to the level of Aβs. Clearly, the conventional BLI strategy is impractical for reporting Aβ levels in vivo. Therefore, innovative tactics are urgently needed. In this report, we hypothesized that AkaLumine, a newly discovered substrate for luciferase ^29^, could bind to Aβ aggregates and plaques that have the reservoir capacity to sequester and release AkaLumine to dynamically control bioluminescence signals in vitro and in vivo (Fig.1). We termed our method as “<u>b</u>ioluminescence imaging with <u>a</u>myloid reservoir (BLIAR)”. For the first time, we demonstrated that, using AkaLumine as the substrate for luciferases, BLIAR could be used to report the levels of Aβ in vivo and monitor the progression of amyloidosis and therapeutic efficacy.

**Fig. 1.**
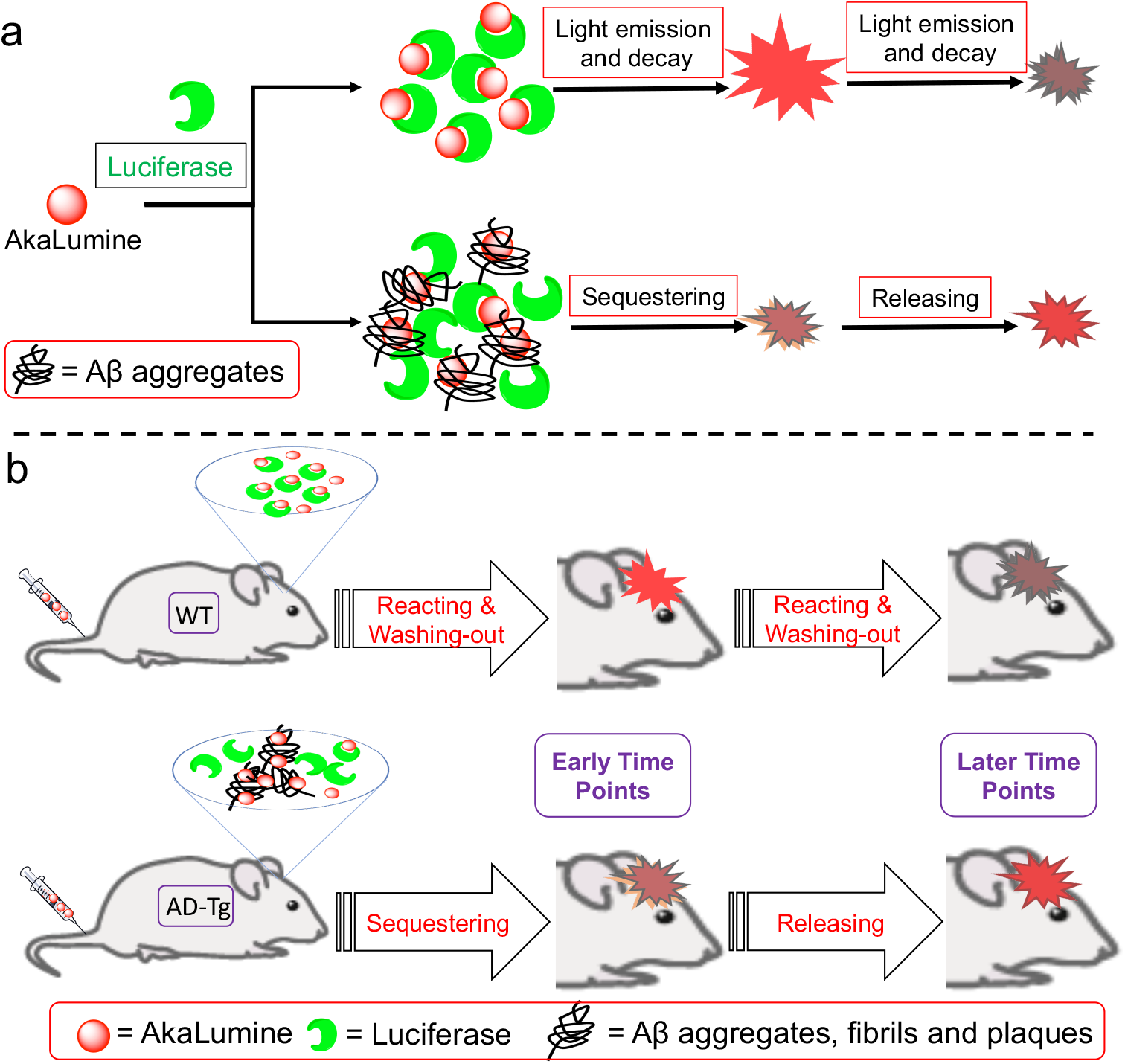
Diagram for illustrating the principle of bioluminescence imaging with amyloid reservoir (BLIAR). a) AkaLumine binds to luciferase to generate bioluminescence and the intensity decrease with time (upper panel); however, in the presence of Aβ aggregates (amyloid reservoir), AkaLumine is first sequestered inside the aggregates and then it is slowly released from the aggregates (lower panel). Compared to the normal bioluminescence generating process (without the reservoir), the bioluminescence intensity from the reservoir group is principally lower at early time points and higher at later time points. b) Illustrating of BLIAR *in vivo*. In normal mice with aka Luciferase (AkaLuc) in brains, AkaLumine reacts with AkaLuc to generate bioluminescence. The bioluminescence intensity decreases with time while AkaLumine is consumed and washed out (upper panel). However, in the brains of AD models, AkaLumine is first sequestered inside the Aβ plaques and then it is released by plaque reservoirs (lower panel). The signals from the AD brains are expected comparably lower at the early time points and higher at the later time points.

## Results

### 1. AkaLumine binds to Aβs

AkaLumine, a newly validated substrate for firefly luciferase (FLuc) and aka Luciferase (AkaLuc), has been demonstrated to have excellent efficiency and unprecedented deep tissue penetration, evidenced by brain imaging of mouse models and marmosets ^29^. Based on its chemical structure, we speculated that AkaLumine could bind to Aβ species, because it shares certain structural similarity with our previously reported Aβ imaging probes CRANAD-Xs (X =-2, -3, -58, etc.) (Fig.2a). In the structure of AkaLumine, the N,N-dimethylamino-phenyl-polyene moiety could be found in CRANAD-Xs and other Aβ-responsive fluorescence dyes (Fig.2a red box and SI Fig.1) ^30-39^. To validate this speculation, we performed competition study between CRANAD-2 and AkaLumine in the presence of Aβ aggregates in PBS buffer (pH 7.4). We found that, indeed, AkaLumine could reduce the fluorescence intensity of CRANAD-2 (Fig.2b). To further confirm that AkaLumine is able to interact with Aβs, we recorded its fluorescence spectra with and without Aβs. We observed an easily discerned intensity increase when Aβ aggregates were added (Fig.2c). Through titration, we calculated Kd value as 37.8 nM, indicating AkaLumine-1 has strong binding to Aβs (Fig.2d). Moreover, for the mixture of AkaLumine and Aβ aggregates, we observed an apparent intensity decrease when the aggregates were removed via spinning down with a centrifuge (Fig.2.e), suggesting that the Aβ aggregates can sequester AkaLumine. In addition, we also performed molecular docking studies based on Aβ40 structures in protein data bank (2MVX), and found that AkaLumine parallelly line with a groove formed by multiple-pieces of Aβ40 peptides. Hydrogen from the carboxylic group in AkaLumine could form hydrogen bond with G37, and hydrophobic interactions could be found between the phenyl ring of F19, L-34 and V-36 (Fig.2f). Taken together, the above results suggested that AkaLumine was able to bind to Aβs in vitro.

**Fig. 2.**
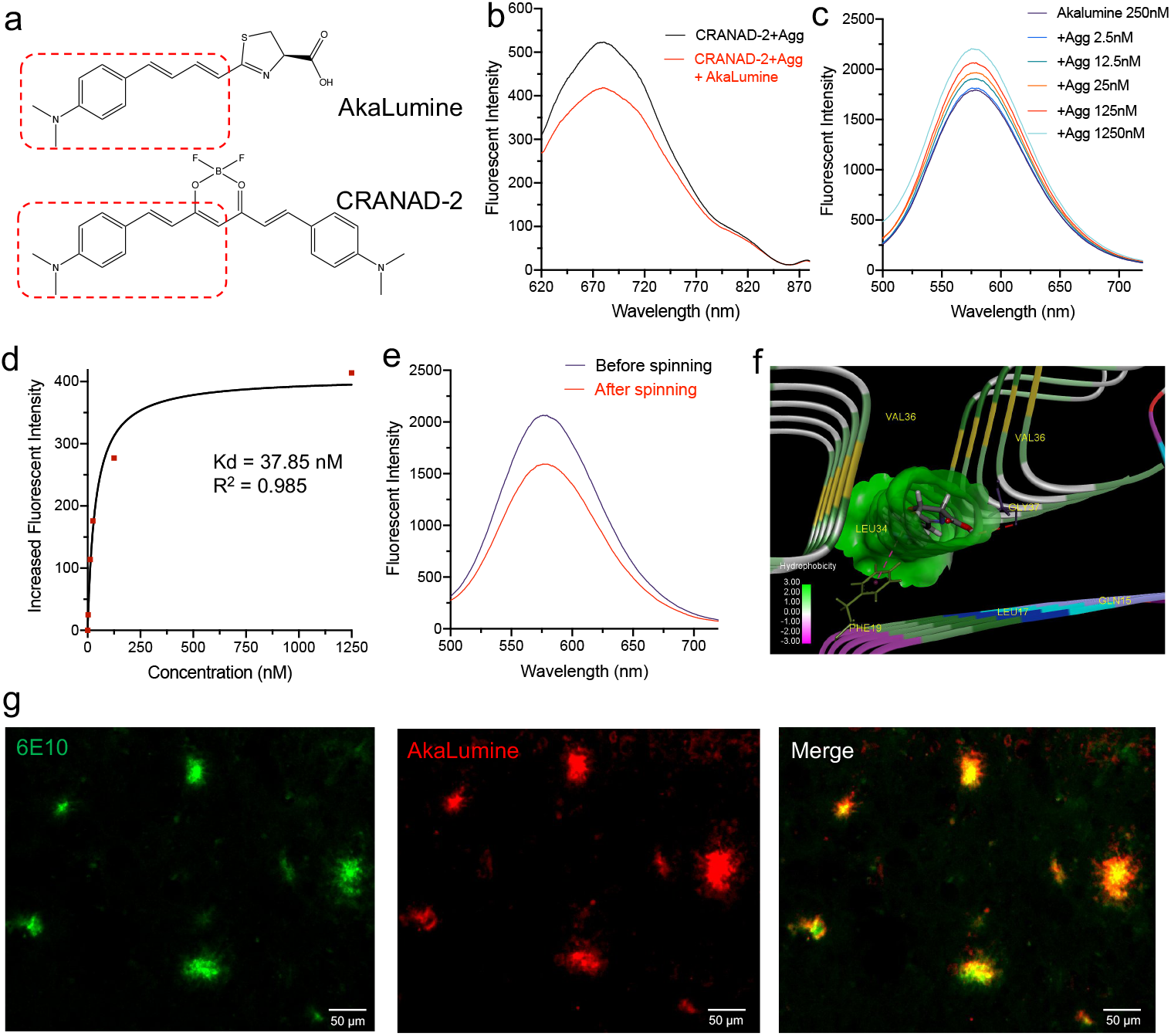
AkaLumine binds to Aβ aggregates and plaques. a) The share structural moiety of AkaLumine and CRANAD-2, a reported fluorescence probe for Aβs. b) Fluorescence spectra of CRANAD-2 to show the competition between CRANAD-2 and AkaLumine in the presence of Aβ aggregates. c) Fluorescence spectra of AkaLumine with different concentrations of Aβ aggregates. d) Binding constant fitting of Akalumine for Aβ binding. e) Fluorescence spectra of the supernatant AkaLumine with Aβ aggregates before and after spinning down with a centrifuge. f) Molecular docking of AkaLumine with Aβ40 fibrils structure (PDB: 2MVX). AkaLumine binds to the hydrophobic pocket formed by Aβ peptides and a hydrogen bond formed between G37 and the hydrogen from carboxylic group of AkaLumine. g) 5xFAD mouse brain slide staining with Aβ antibody 6E10 (left, green) and AkaLumine (middle, red). The staining from AkaLumine and 6E10 showed excellent co-localization (right, merged image).

To further investigate whether AkaLumine can bind to Aβs in a biologically relevant environment, we stained brain slides from an AD mouse, and found that it could provide excellent labelling for Aβ plaques (Fig.2g). The labeled plaques were further confirmed by immunohistology staining with 6E10 antibody, a specific antibody for Aβs ^40^ (Fig.2h,i). Interestingly, we found that firefly luciferin was not able to stain the plaques (SI Fig.2).

### 2. Aβ fibrils/aggregates and plaques are reservoirs for AkaLumine

Given that Aβ fibrils/aggregates contain numerous binding sites that are created by thousands of beta-sheets of Aβ peptides, we hypothesized that Aβ aggregates could be considered as reservoirs to temporarily sequester and later release AkaLumine, which could dynamically control the availability of AkaLumine to luciferase to generate bioluminescence (Fig.1). Based on this hypothesis, we reasoned that at the beginning period, the bioluminescence intensity in the presence of Aβ aggregates will be lower than that from the control group (without Aβ aggregates); however, at the later time-points, there will be a reversed trend of the intensity, i.e. the intensity from the Aβ group will be higher than that from the control group. To validate this hypothesis, we performed imaging experiments with Eppendorf tubes with filter inserts (Fig.3b). Since AkaLuc is not commercially available, we used FLuc, which also can catalyze AkaLumine to produce bioluminescence, evidenced by the fact that the peak 675nm of bioluminescence produced by FLuc with AkaLumine is consistent with the reported value ^29,41^ (SI Fig.3). In the experiments, we first added AkaLumine/FLuc/ATP or AkaLumine/FLuc/ATP/Aβ aggregates to the inserts, and then imaged the inserts with an IVIS imaging system. As expected, we observed that the intensities from the Aβ inserts were significantly lower than that from the control inserts (no Aβs) (Fig.3a).

**Fig. 3.**
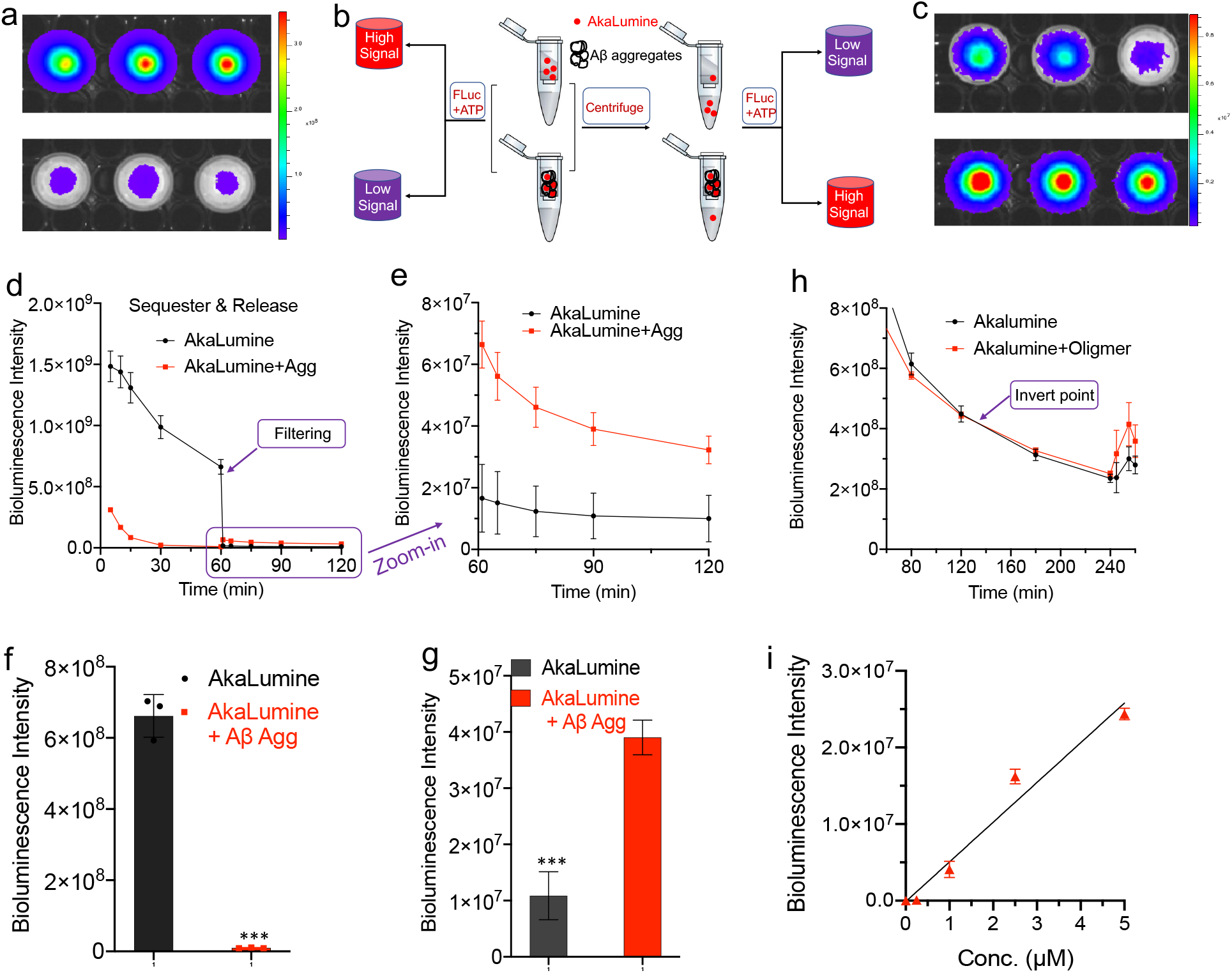
Validation of Aβ aggregates as reservoirs to sequester and release AkaLumine in solutions. a-c) Higher signals were observed from the group without Aβ aggregates (a) based on the experimental procedure outlined in (b), and reversely lower signals were observed from the same control group (without Aβ aggregates) after filtering (c). d-e) Quantitative analysis and time course of bioluminescence intensities before filtration (d) and after filtration (e). Inverted intensities can be easily observed from the control group (black lines) and the Aβ group (red lines). f-g) The largest margins from the two groups before filtration (f) and after filtration (g). h) Partial time course of bioluminescence intensities from AkaLumine with and without Aβ oligomers. An inverted point can be seen between 120 to 180 minutes. i) Linear fitting of Aβ aggregates concentrations under 5.0 μM. Error bar: Stdev. P value, ** < 0.01, *** < 0.001.

The apparent differences could be observed for all the time points (Fig.3d), and the largest difference reached 70-fold (control/ Aβ) at 60-minutes after the starting of the reaction (Fig.3f). To mimic the washing-out effect in vivo, we spun the inserts with a centrifuge to filter away free AkaLumine. After the spinning, solutions of FLuc/ATP were re-added to all the inserts, which were subjected to re-imaging. Indeed, the intensity trend was inverted after the filtering, evidenced by the higher intensity from the Aβ group (Fig.3c,e,g). The difference reached 3.71-fold (Aβ/control) at 30 minutes after the re-adding of FLuc/ATP (Fig.3g). Our results suggested that Aβ aggregates could have the reservoir function to store and release AkaLumine.

Moreover, we also observed similar phenomena with Aβ oligomers, indicating our method is not only suitable for insoluble Aβs, but also for soluble Aβs (Fig.3h and full time-course in SI Fig.4), which is believed to be the early-stage biomarkers of AD. Interestingly, the inverting point could be observed in Fig.3h around 120 minutes without filtering; however, the larger difference could be found after washing away free AkaLumine through filtering (Fig.3h). This margin of difference from the case of Aβ oligomers is much smaller than that from aggregates (Fig.3d *vs* Fig.3h), presumably due to less trapping capacity of Aβ oligomers. In addition, we found that the reduction caused by Aβ aggregates was linear to its concentration in a range of < 5.0 μM (Fig.3i).

### 3. In vitro and in vivo mimic studies

We next used brain homogenates as in vitro mimic conditions to verify the sequester-release hypothesis. In this regard, we incubated AkaLumine (100 nM) with WT and 5xFAD brain homogenates for 20 minutes under shaking. After the incubation, the homogenates were centrifuged, and the pellets were collected. Half of the pellet was subjected to extraction with organic solvent ethyl acetate. We measured the fluorescence of the extractions with a fluorometer. Indeed, we found that the intensity from the 5xFAD group was 2.47-folder higher than that from the WT group (Fig.4a), suggesting that Aβ plaques in the homogenate have the sequestering capacity. Another half of the pellet was resuspended with PBS buffer. After a certain period, the suspension was centrifuged, and a portion of the supernatant was taken. We attempted to measure the fluorescence of the supernatant, but we found the signal is weak. However, we were successful to measure AkaLumine fluorescence after extracting with ethyl acetate. This is because most of fluorescent compounds have much stronger fluorescence in organic solvents, such as ethyl acetate and dichloromethane. Indeed, we found that the pellets could slowly release AkaLumine, evident by the increased fluorescence of AkaLumine in the PBS supernatant with the resuspending time. The intensity at 5-hours was 1.28-fold higher than that at 0-minute in the 5xFAD group, while only 1.09-fold higher intensity was observed in the WT group during the same period (Fig.4b). Taken together, our data support our sequester-release hypothesis in brain tissues.

**Fig. 4.**
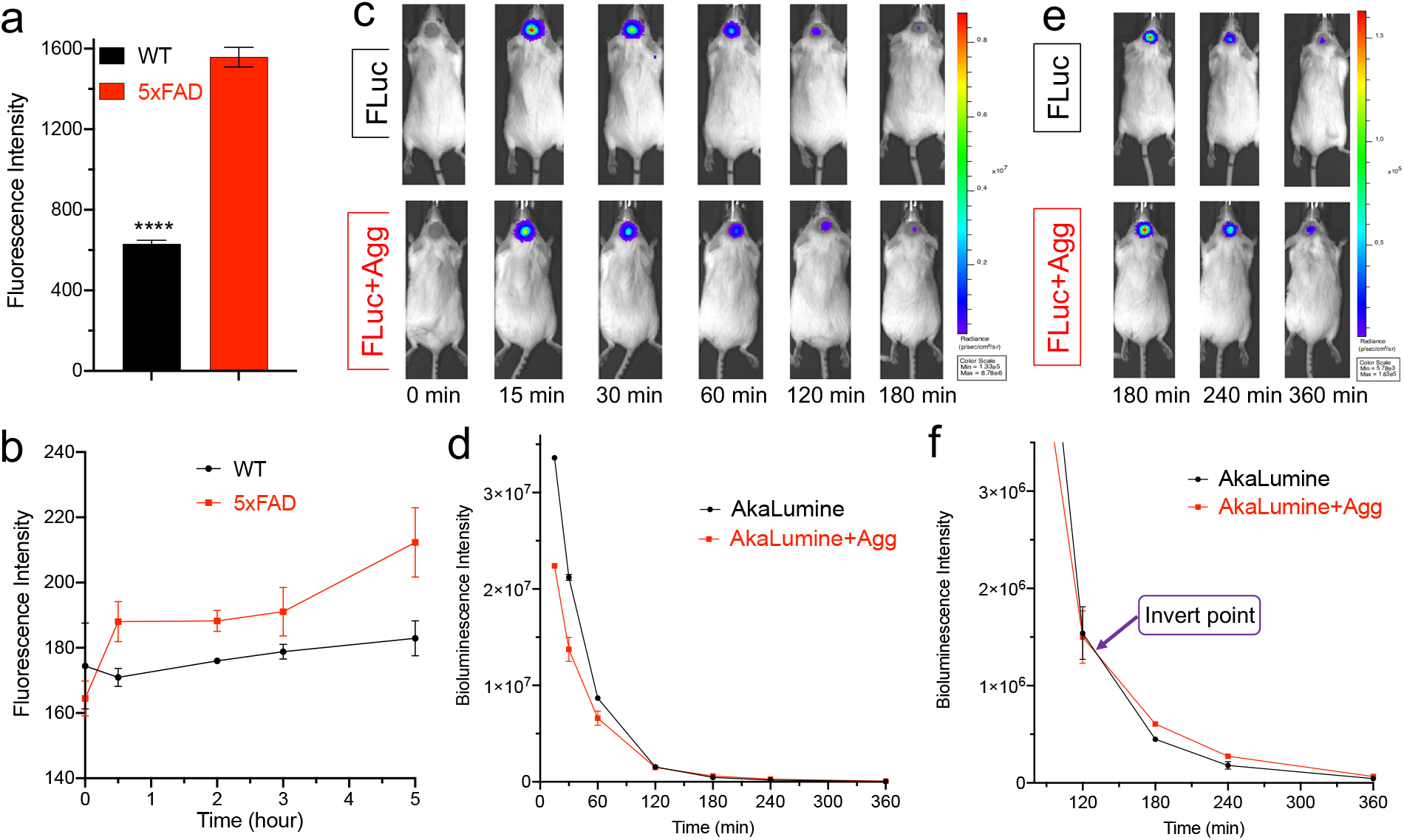
In vitro and in vivo mimic studies for validating the reservoir function of Aβs. a-b) In vitro brain tissue homogenates for validating sequester-release of AkaLumine. Fluorescence intensities of AkaLumine from pellets of brain homogenates (WT and 5xFAD) that were incubated with AkaLumine (n=3). b) Fluorescence intensities of the released AkaLumine at different time-points from the pellets that were resuspended with PBS solutions (n=3). c) In vivo bioluminescence imaging at early time points (0-180 minutes) in mice that were subcutaneously inoculated with FLuc only (upper panel) and Aβ aggregates + FLuc (lower panel) under the scalp. d) Quantitative analysis of images for the full time-course (0-360 minutes) after iv injection of AkaLumine (n = 2). e) In vivo bioluminescence imaging at later time points (90-360 minutes) in mice that were subcutaneously inoculated with FLuc only (upper panel) and Aβ aggregates + FLuc (lower panel) under the scalp. f) Quantitative analysis of images for later time-points (90-360 minutes) after iv injection of AkaLumine (n = 2). An inverted point can be clearly seen around 120 minutes after AkaLumine injection. P Value: **** < 0.0001. Error bar: Stdev.

To investigate whether our hypothesis is achievable under in vivo conditions, we first performed in vivo BLIAR mimic experiments, in which we performed subcutaneous inoculation of Fluc or Fluc+ Aβs under the scalp and then intravenous injection of AkaLumine. Indeed, we observed 1.68-fold higher BLI signals from the control group, and the difference margin was gradually becoming smaller from 0 min to the inverting time-point (Fig.4c,d). As expected, a reversing point could be seen around 120 min after the injection of AkaLumine. The intensity ratio of Aβs/control groups could reach 1.96-fold at 360 min (Fig.4e,f). Our data indicated that Aβs could also have the reservoir function under the in vivo mimic conditions. To further confirm our hypothesis, we performed in vivo mimic ocular imaging, in which FLuc or FLuc + Aβs was injected into eyes and AkaLumine was then intravenously injected. As expected, the signals from the Fluc only group are significantly higher than that from the Fluc + Aβs group, and the difference was 4.61-fold at 60 minutes after AkaLumine injection (SI Fig.5a,b), suggesting that our method is potentially feasible under in vivo conditions. However, we did not observe an inverting point, because the signal was quite low after several hours.

### 4. In vivo two-photon imaging and evidence of sequestering-releasing

For in vivo brain imaging, penetration of blood brain barrier (BBB) and labeling plaques are the premises for an AD imaging probe. In this regard, we examined whether AkaLumine could cross BBB and label Aβ in vivo. Using two-photon imaging with a 5xFAD mouse, we confirmed that AkaLumine could cross BBB and provide excellent contrast for Aβ plaques and cerebral amyloid angiopathy (CAA) on the blood vessels (Fig.5a,b). The capacity of AkaLumine for labeling Aβ plaques and CAAs is consistent with our results from in vitro brain slide staining (Fig.2g).

**Fig. 5.**
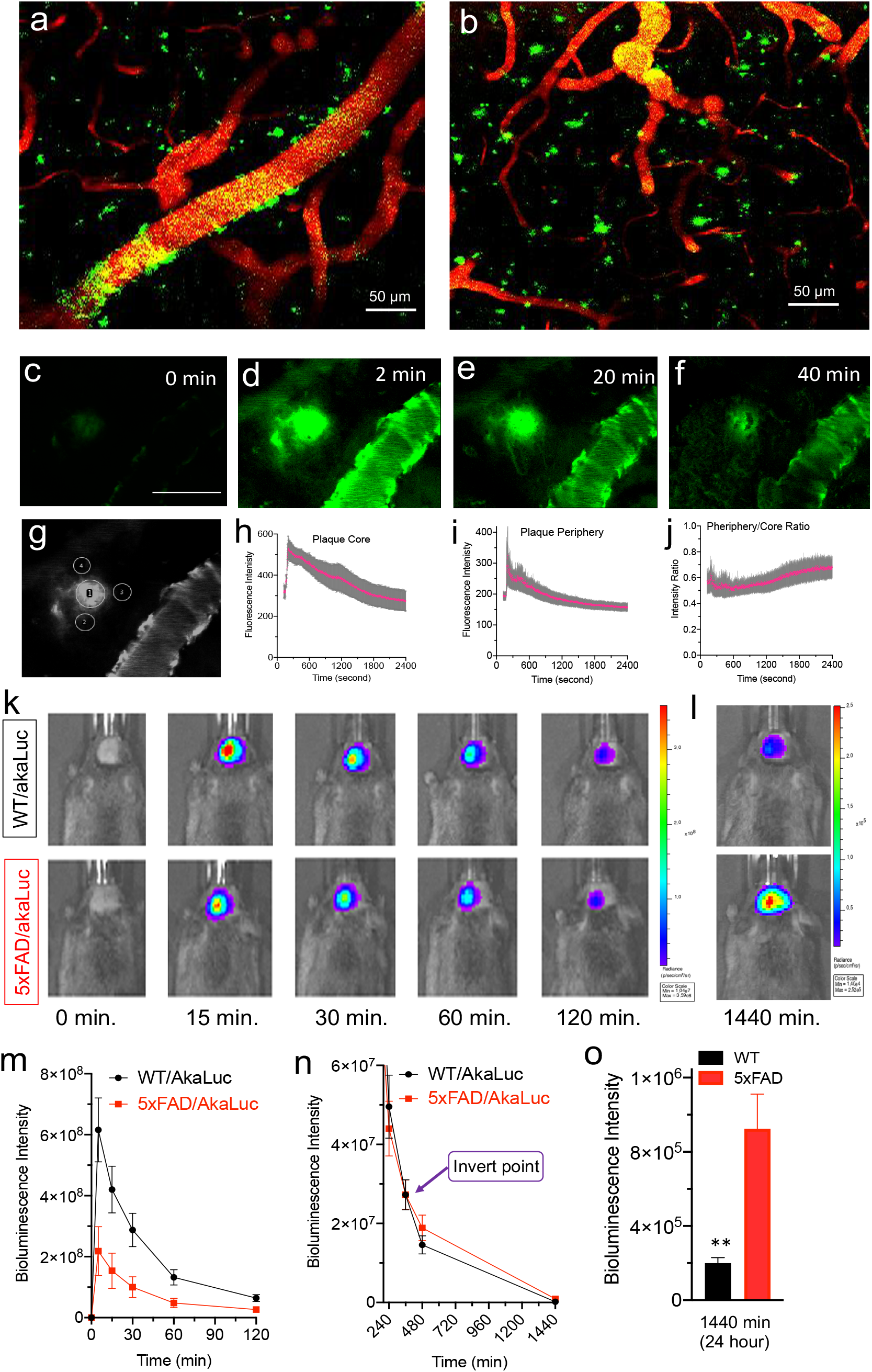
In vivo two-photon microscopic fluorescence imaging (a-j) and in vivo bioluminescence imaging (k-p) with AkaLumine. a) Cerebral amyloid angiopathy (CAA) Aβ deposits (green) labelled by AkaLumine on blood vessels (red) in the brain of a 12-month-old 5xFAD mouse. b) Aβ plaques (green) labelled by AkaLumine in a cortex area. c-f) Representative images of a plaque at different time points after AkaLumine injection (scale bar: 50 micron). g) Representative region of interest (ROI) for quantification. h) Time-course of plaques (n=6). i) Time-course of peripheral areas of plaques (n=18). j) Time-course of the intensity ratio of periphery/core. Error bars: SEM. k) In vivo bioluminescence imaging at early time points (0-120 minutes) with WT/AkaLuc (upper panel) and 5xFAD/AkaLuc (lower panel) (n=4). Apparently higher signals can be observed from the WT/AkaLuc group. l) Representative images at a later time point (1440 minutes/24 hours) of WT/AkaLuc and 5xFAD/AkaLuc groups. Apparently lower signals can be observed from the same WT/AkaLuc group (upper panel). m) Time-course of bioluminescence intensities for the early time points (0-120 minutes). n) Time-course of bioluminescence intensities for the later time points (240 -1440 minutes). o) Quantitative analysis of images at 24-hours for WT/AkaLuc and 5xFAD/AkaLuc groups. Bioluminescence intensity from the 5xFAD/AkaLuc group was 4.26-fold higher than that from the WT/AkaLuc group at this time point. P value, ** < 0.01, *** < 0.001.

To investigate whether plaques have sequestering-releasing capacity, we acquired time-lapse images for 40 minutes after AkaLumine injection. We found that, indeed, plaques could quickly sequester AkaLumine after injection, and the fluorescence intensity of plaques reached the peaks around 2 minutes post injection (Fig.5c,d,h). After that, the plaques slowly released AkaLumine, evident by the loss of contrast between the core and periphery of a plaque (Fig.5e,f). To further confirm the sequestering-releasing capacity of plaques, we quantified individual plaque and its three peripheral areas (Fig.5g), and time-courses were plotted (Fig.5h, I,j). Although the intensities both from the core and the periphery were decreased, due to the probe washing-out effect, the rate of the decrease from the cores were faster than that of the peripheral areas. Consistent with this phenomenon, it was clearly that the intensity ratio of periphery/core was slowly increasing during the acquisition of images (Fig.5j), suggesting that the sequestered AkaLumine in the core was slowly released to the peripheral areas. Similarly, we have observed similar trends from CAAs (SI Fig.6a-c). Taken together, our data support the sequestering-releasing hypothesis.

### 5. In vivo reporting Aβ levels via BLIAR with a transgenic AD mouse model

To investigate whether our hypothesis is applicable in transgenic AD mouse model, we have created two strains of mice via intracerebral viral injection of AAV2 with packed DNA of AkaLuc. Although FLuc is effective enough for in vitro and mimic in vivo studies, it is not the best matched enzyme for AkaLumine. In 2018, Iwano et al optimized luciferase via mutagenesis, and found that AkaLuc was the best match for AkaLumine to generate bioluminescence in the near infrared window ^29^. In this regard, for in vivo brain imaging, we selected AkaLuc to match with AkaLumine ^29^. To this end, we constructed plasmid DNA (pAAV-SV40LpA-tTAad-SYN-insulator-TRE-Venus-AkaLuc-BGHpA) based on the reported protocol, which was subsequently packaged with AAV2 ^29^. Ten-month old 5xFAD mice and aged-matched wild type mice were microinjected with AAV in the cerebral cortex under direct visualization through a temporary cranial window. Three weeks were allowed for wound healing and virus expression.

First, we validated that the AkaLuc/AkaLumine pair could provide longer emission, evidenced by the 670nm peak of the emission spectrum from in vivo brain imaging (SI Fig.6d). In this study, the wild type mice with AAV viral injection (WT/AkaLuc) were used as the control group, while the 5xFAD mice with AAV viral injection (5xFAD/AkaLuc) were used as the experimental group. Based on our hypothesis, we expected that the bioluminescence signal from the 5xFAD/AkaLuc group would be much lower than that from the control group at the early time-points, because the Aβ plaques can sequester the injected AkaLumine. Indeed, we found that the signals from the 5xFAD/AkaLuc group was 2.8-fold lower, compared to the control group (Fig.5c,e), and the difference lasted for almost 4 hours (Fig.5e,f). Expectedly, the difference become smaller with time (Fig.5c,e), and the signal decay was faster in the control group (4.73 vs 2.79 of the decay rate). Our in vitro solution tests showed that an inversion trend of difference could observed after filtering (Fig.3a-g). Similarly, we observed the inversion in vivo around 360 minutes post AkaLumine injection (Fig.5d,f,g), and the ratio of 5xFAD/AkaLuc and WT/AkaLuc could reach 4.64-fold at 24-hours after AkaLumine injection (Fig.5d,g). Compared to the control group, the lower signals at early time points and higher signals at later time points make our imaging method unique. With one set of experiments, we were able to obtain two data sets (the BLI signals before and after inversion), and these data series can be used to cross-validate each other. This will significantly reduce artifacts to make the results more reliable. Taken together, our results clearly validated our hypothesis.

To rule out different expression levels of AkaLuc in WT and 5xFAD groups, we sacrificed the mice and performed ex vivo brain imaging. Since DNA of Venus was included in our plasmid construct, we used the ex vivo brain imaging to quantify the fluorescence intensity from WT/AkaLuc and 5xFAD/AkaLuc, and found that the two groups had similar intensities (SI Fig.7), further confirming that the expression levels of AkaLuc were similar.

### 6. BLIAR for monitoring progression and therapeutic effectiveness

For preclinical animal cancer research, molecular imaging, particularly bioluminescence imaging, has been routinely used to monitor the effectiveness of drug treatment and genetic manipulation. However, it is still scarce to use molecular imaging methods to monitor treatment efficacy in AD preclinical animal research. To investigate whether BLIAR can monitor changes of Aβ burden during ageing and drug treatment, we first used different ages of 5xFAD/AkaLuc mice to perform BLI imaging at 5 minutes post i.v. injection of AkaLumine. As expected, at the age of 8-month-old, 1.78-fold lower BLI signal was observed from the 5xFAD/AkaLuc group (n = 4), compared to the WT/AkaLuc group (n = 4) (Fig.6a). Similarly, the BLI signal was 3.48-fold lower in 5xFAD/AkaLuc group (n = 6) than that in WT/AkaLuc group (n = 6) at the age of 12-month-old (Fig.6b). However, we found that the difference between the 8-month-old groups was not significant (p = 0.0768). This is likely due to the variations of AkaLuc expression level in individual mouse. To minimize the effect of varied expression, the decay of BLI signal was fit with two-phase decay model, and we quantified the halftime (T1/2) to further validate the above results (SI Fig.8a,b). We found that there were no significant differences of the T1/2 of the fast phase between the 5xFAD and WT groups (SI Fig.8e). This is reasonable because the fast phase is likely dominated by non-bound AkaLumine, which is faster to react with AkaLuc than the bound AkaLumine in the plaques. Expectedly, the T1/2 of slow phase from both 8- and 12-month-old 5xFAD/AkaLuc mice were significantly different, compared to the WT/AkaLuc mice. Moreover, the T1/2 of 12-month-old 5xFAD/AkaLuc mice was 1.27-fold longer than that of the 8-month-old 5xFAD/AkaLuc mice (Fig.6c); whereas there was no difference of T1/2 between the WT groups. Taken together, our data suggested that BLIAR could be used to monitor the progress of amyloidosis in transgenic AD mice.

To further demonstrate that BLIAR can be used for monitoring the changes of Aβ burden in vivo, we treated 8-month-old 5xFAD/AkaLuc mice with gamma-secretase inhibitor avagacestat, which has been well-documented for its efficacy to reduce Aβ burden in mouse models, dogs and human subjects ^42-44^. After two months of treatment (i.p. three times/week), we imaged the mice with i.p injection of AkaLumine. After imaging, the mice were sacrificed and brains were imaged with Venus fluorescence, which can be used to reflect the expression level of AkaLuc. With normalization of Venus intensity, we found that the BLI signal from the treated group (n = 3) was 2.38-fold higher than that from the non-treated control group (n = 3) at 240 minutes after AkaLumine injection (Fig.6d), and the differences could be observed through the time course (Fig.6e). We also compared the trajectory of individual mouse before and after drug treatment and vehicle treatment. We found that the changes were slower in the avagacestat treated group than that in the vehicle treated group (Fig.6f), suggesting drug treatment could slow down the of plaque growth and accumulation. We further validated the reduction of Aβ burden with immunoassay (Mesoscale detection, MSD), and found that insoluble Aβ42 concentration was 1.82-fold lower in the treated group, compared to the control group (Fig.6g). Similarly, 1.20-fold lower insoluble Aβ40 burden was detected from the treated group (SI Fig.9).

**Fig. 6.**
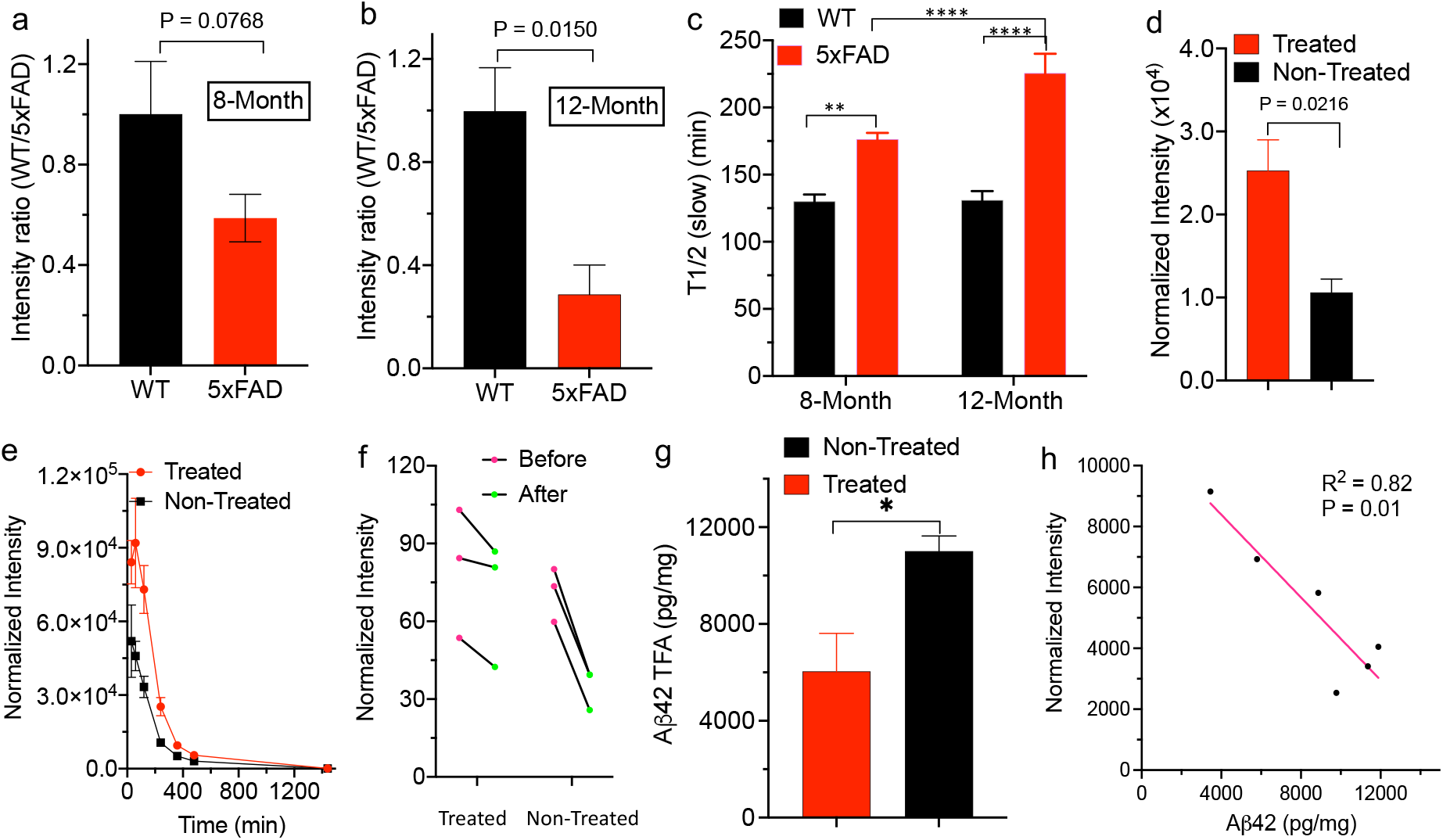
Monitoring AD progression and therapeutic effectiveness with BLIAR. (a,b) BLI intensity differences at 8- and 12-month of WT/AkaLuc and 5xFAD/AkaLuc mice (n=4). (c) Half-time (T1/2) of WT/AkaLuc and AD/AkaLuc mice at 8- and 12-month by two-phase decay fitting (slow phase). Both BLI intensity and half-time results indicated that BLIAR could be used to monitor the increase of Aβ burden in vivo. (d) Normalized BLI intensity showed significant difference between avagacestat-treated and non-treated group at 240-minutes after two-months therapy. (e) Time-courses of the treated and non-treated groups also showed consistent differences at different time points. (f) The trajectory of individual mouse before (red) and after (green) drug or vehicle treatment. (g) ELISA analysis with MSD Aβ kit for brains from the treated and non-treated group. The treated group showed lower Aβ42 levels in the treated group. (h) Inverse correlation between BLI and Aβ42. BLIAR signals showed excellent correlations with insoluble Aβ42 concentrations. P value: * < 0.05, ** < 0.01, *** < 0.001.

To evaluate whether an imaging method is robust enough to monitor changes from pharmacological and genetic manipulation, an excellent linear correlation between imaging signal and biomarker level is required. In this regard, with the treatment study, we performed linear regression fitting with BLI signals and Aβ42 concentrations from MSD measurement. Indeed, we found that excellent linear correlations for BLI signals at 240-, 480- and 1440-minutes (Fig.6h and SI Fig.9). The R^2^ value was 0.82 and p value was 0.01. As expected, the relationships are negative correlations. Collectively, our results indicated that BLIAR could be used to monitor the therapeutic effectiveness in mouse models.

## Discussion

BLI has been rarely used in AD mouse models ^25-28^. Watts et al reported that BLI could be used for AD mouse imaging; however, the FLuc was engineered with promoters of Gfap and mutant human amyloid precursor protein (APP) ^25^, which primarily reflected the level of GFAP and astrocyte activation in the progress of AD pathology, and the correlation between the GFAP/APP level and the deposit of Aβs. Similarly, Dunn-Meynell used BLI for qualifying GFAP with Gfap-FLuc engineered mouse model for studying Tau pathology. Although correlations between Aβ deposits (and/or tau tangles) and GFAP levels could be observed; however, strictly the level of GFAP could not be used to report the level of Aβ deposits. Van der Jeugd et al co-expressed luciferase with human hTau40/ΔK280 in mice, and achieved stable BLI signal at the age of > 4-months. However, they also reported an apparent drop of BLI signals from 1-month old to 4-month-old (3-fold decrease) ^11,24^.

To the best of our knowledge, our report is the first of its kind of in vivo bioluminescence imaging that can be used to report the level of Aβs and changes of Aβ burden, which is desperately needed for the community of preclinical AD research. Compared to previously reported fluorescence imaging ^30-39^, our method also provided considerable larger difference between AD mice and control mice, and this is very crucial for monitoring disease progression and effectiveness of drug treatment. Conceivably, our strategy can be extended to other misfolding-prone proteins, including tau proteins, synucleins, SOD, TDP-43 etc ^45^. Conjugation of AkaLumine with a ligand of these proteins or modifying AkaLumine to make its analogues as their ligands are plausible approaches.

Compared to fluorescence imaging, BLI is much more straightforward and easier to operate, and always user friendly, and more importantly, BLI normally provides considerably high signal-to-noise ratios (>1000) ^19^, which are essential for reliable quantification. BLI has been widely applied for various purposes; however, most of studies of in vivo whole-body fluorescence imaging are still at the stage of probe validation and few probes have been widely applied, due to different fluorescence probes have different excitation and emission, which requires expertise to properly setup the experiments. Additionally, the low signal-to-noise ratios of NIR fluorescence imaging has significantly hampered its wide applications ^19^. Contrary to in vivo fluorescence imaging, BLI has significantly promoted the progress of drug development at preclinical stages for various diseases, particularly for cancer research. In the past three decades, BLI has profoundly changed the daily practice for cancer research ^21,22^. Nonetheless, BLI has been rarely used in preclinical AD studies to monitor the progress of disease ^11,24-27^. With our method, the day of such change is possible to come. Given that BLI can be performed easily by non-imaging experts, our BLIAR method has paved the road for widely using BLI for Aβ level monitoring for both preclinical and basic mechanism studies. We believe that our BLIAR has great potential to change the daily practice of AD research in both academic and industrial laboratories.

In vitro tissue studies suggested that more AkaLumine was retained in AD brain homogenates than that in WT, and the homogenate pellets could slowly release AkaLumine. Moreover, our two-photon imaging studies not only confirmed that AkaLumine could bind to plaques and CAAs in vivo, but also provided direct evidence to support our sequestering-releasing hypothesis. However, it is impracticable to continuously track the intensity changes of a plaque and its peripheral areas for long periods of time (> 60 minutes), due to the possible photo- and heat-damage to the imaging areas with long-time laser irradiation.

Numerous studies suggest that Aβ plaques are not static, but rather they grow and expand dynamically throughout AD progression. Hyman et al show that newly formed plaques can alter the curvature of neurites ^46^, and a profound deficit in spine density is often observed around plaques ^47^. Bacskai et al show that the plaques inhibit mitochondrial function and calcium homeostasis, and lead to rapid cell death in the vicinity of the plaques due to oxidative stress ^48,49^. Condello et al show that plaques grow gradually over months with growth slowing in older AD mice, and the degree of neuritic dystrophy correlates with the speed and extent of plaque enlargement. Remarkably, new plaques induce a disproportionately large area of neuritic dystrophy ^50 51^. We also suggested that the plaques could be divided into “active” and “silent” plaques ^52^. Due to the different states of plaques, at the micro-level, AkaLumine could have different pharmacodynamics (PD) and pharmacokinetics (PK) in each category of plaques. At the macro-level, the dynamic profile of bioluminescence signals from AkaLumine could be different in AD mice at different ages. We will perform more detailed studies in the future to investigate the correlations between the PK/PD at micro-level and macro-levels in AD mice.

It has been reported that brain blood barrier (BBB) of AD mice could be disrupted, and the CAA deposit could have impact on the influx and exflux of polar small molecules and large molecules ^53,54^. Since AkaLumine is a non-polar small molecule and its molecular weight < 300 da, it is likely that it could freely cross BBB, and it is reasonable to believe that CAA has no significant effect on AkaLumine’s BBB penetration. In addition, both in vitro slide staining and in vivo two-photon imaging indicated that AkaLumine had no strong binding to microglia and astrocytes, evident by the lack of microglia and astrocyte morphology from the images. However, more studies are needed to confirm this preliminary observation ^55^.

Fluc and firefly luciferin is the most used pair for BLI, and we have attempted to investigate whether this pair can be used for our purpose. However, we found that luciferin was not able to label Aβ plaques of AD mouse brain slides (SI Fig.2). In addition, compared to AkaLuc/AkaLumine pair, the FLuc/Luciferin pair provides much shorter emission wavelength, which further limits the application of this pair for brain imaging. AkaLuc/AkaLumine is the ideal pair for in vitro studies; however, we used FLuc to replace AkaLuc for in vitro studies, due to the commercial unavailability of AkaLuc. We did not foresee that using FLuc could comprise our conclusions of in vitro studies, because FLuc is also capable to bind to AkaLumine to produce excellent BLI signals.

For our in vivo imaging, instead of using Fluc as the enzyme, our transgenic AD mice were engineered with the best matched AkaLuc enzyme via viral transduction of AkaLuc DNA in the cerebral cortex area. As we expected, the brain imaging results are robust and very supportive for hypothesis. However, local intracerebral AAV viral transduction is an invasive procedure and brings considerable expression variations for individual injection. To avoid these problems, in the next step, we plan to create mouse models that have both AkaLuc and AD genes (such as APP, PS1) specifically expressed in brains. For example, Rosa26-Syn-AkaLuc strain of mice can be specifically knocked-in for a brain and sub-areas of a brain ^56^, and cross-breeding of this strain with transgenic AD mice (APP/PS1, 5xFAD) can produce strains of AD/AkaLuc mice. In 2018, Iwano et al showed that the AkaLuc/AkaLumine pair could be used for brain function imaging in marmosets ^29^. We envision that this is feasible for marmoset AD models as well, which may have much better capacity to recapture human AD features. Moreover, we speculate that BLIAR is feasible to report the Aβ levels in eyes in transgenic mice and large animals such as beagle dogs and marmosets, which can be locally engineered with AkaLuc in eyes. In addition, utilizing our method for future clinical studies is potentially viable via ocular imaging. Conceivably, it is practical to locally inject luciferase (Fluc or AkaLuc) and AkaLumine into eyes, and the level of ocular Aβs can be measured via BLI quantification.

In this report, we showed that Aβ plaques could lead to signal reduction at early time points, and we used to WT/AkaLuc as the control to relatively quantify the levels of Aβs to validate our hypothesis. However, in daily practice, the WT/AkaLuc mice may not need in the course of investigating the changes of Aβ levels after drug treatment or genetic or pharmacological manipulation. Reasonably, if the treatment or manipulation can reduce the Aβ levels, we anticipate BLI signal increases at the early time points while decreases at later time-points after the intervention. Indeed, this speculation is consistent with our treatment study with avegacestat. Remarkably, in this treatment study, we revealed excellent inverse correlations between BLI signals and Aβ levels that were measured by MSD analysis of brain homogenates. The excellent correlations strongly suggested that BLI signals could be used to report the Aβ levels in vivo.

In this report, we demonstrated that BLIAR could be used for real-time monitoring Aβ burden changes during disease progression and after treatment with gamma-secretase inhibitor avegacestat, which has failed in clinical trials but has well-validated Aβ reduction efficacy across species, including mouse, dog and human ^42-44^. Several groups are working on gamma-secretase modulators, which have potential to avoid the problem of inhibition of Notch cleavage that are associated with the gamma-secretase inhibitors ^57,58^. The treatment study was preliminary, due to small sample sizes of the drug-treated and vehicle injection groups. Large cohorts with younger mice are needed in the future to further confirm the monitoring capacity of BLIAR. In addition, we found that AkaLuc expression via viral transduction in some batch injections had small variations, but other batches had larger variations. However, this problem could be partially solved using transgenic AD mice that has stable expression of AkaLuc in brains. This work is currently ongoing in our laboratory via collaborative efforts. In addition, optimizing studies are needed to establish standard operation procedures (SOP) for routine applications once the transgenic AD/AkaLuc mouse model is ready for use.

Currently, to assess the levels of Aβs after drug treatment or genetic and pharmacological manipulation, ELISA, plaque number and area counting are the most used methods. However, these methods are expensive, due to the need to sacrifice the expensive AD mice for each manipulation or each time point. Moreover, those methods can’t be used for real-time monitoring, as they are ex vivo tissue based. In addition, those tissue-based methods are error-prone, due to each step, such as homogenizing, extraction, antibody incubation, and washing, could lead to errors.

Although a few examples of considering Aβ species as reservoir have been reported ^59-62^, none of them has considered this reservoir can be used to sequester small molecules and later release them. In all those cases, the reservoir capacity has been investigated as self-dynamics of the Aβ monomers in and out of the aggregates or plaques. Our hypothesis is unique and has change the way to utilize Aβ plaques. Prospectively, based on the results of our studies, it is possible to utilize Aβ plaques to control release small drug molecules that can tamper down neurotoxicity of the plaques, inflammation and/or the over-activation of immune activity around the plaques.

In summary, in this report, we demonstrated the feasibility of bioluminescence imaging of Aβ species and monitoring changes of Aβ burden in vivo. Our method can be easily adapted by regular biology laboratories. We believe that this method has the potential to change the daily practice of preclinical AD research and will greatly assist AD drug discovery and development.

## Materials and Methods

All the chemicals were purchased from commercial vendors and used without further purification. AkaLumine (TokeOni) was purchased from Sigma Aldrich (Cat. No. 808350); Aβ1-40 HFIP (A-1153-2) and Aβ1-40 TFA (A-1001-2) were purchased from rPeptide; Anti-β-amyloid antibody 6E10 was purchased from Biolegend (Cat. No. 803001); pcDNA3 Venus-AkaLuc was purchased from RIKEN BioResource Research Center (Clone ID: RBD-15781). pcDNA3 Venus-AkaLuc plasmid was packed with AAV2 virus and purified by Vigene Biosciences. All animal experiments were approved by the Institutional Animal Use and Care Committee at Massachusetts General Hospital.

### Preparation of Aβ40 aggregates

1.0 mg of Aβ40 peptide (TFA) was suspended in a 1% ammonia hydroxyl solution (1.0 mL), then 100.0 μL of the resulting solution was taken and diluted 10-fold with PBS buffer (pH 7.4) and stirred at room temperature for 3 days. Transmission electron microscopy and Thioflavin T solution test were used to confirm the formation of aggregates ^30^.

### Preparation of Aβ40 oligomers

1.0 mg of Aβ40 peptide (HFIP) was suspended in 1.0 mL HFIP, then 20.0 μL of the resulting solution was diluted 10-fold with PBS buffer (pH 7.4) in an eppendorf tube, which was left open at room temperature overnight till HFIP was completely volatilized. To this tube, PBS buffer (20.0 μL) was then added to make a solution of 200.0 μL. Transmission electron microscopy were used to confirm the formation of oligomers.

### In vitro fluorescence spectral test in solutions

Solutions of AkaLumine (250 nM) in 1.0 mL PBS, or AkaLumine (250 nM) with Aβ aggregates (2.5nM, 12.5 nM, 25 nM, 125 nM and 1250nM) in 1.0 mL PBS were prepared, and fluorescence spectra of the prepared solutions were recorded with excitation at 380 nm and emission from 500 nm to 750 nm using an F-7100 fluorescence spectrophotometer (Hitachi). A blank control of PBS was used to correct the final spectra. For CRANAD-2 competition study, solutions of CRANAD-2 (250 nM) with Aβ aggregates (250 nM) in 1.0 mL PBS, or CRANAD-2 (250 nM) with Aβ aggregates (250 nM) and AkaLumine (125 nM) in 1.0 mL PBS were prepared, and the fluorescence spectra of the prepared solutions were recorded with excitation at 580 nm and emission from 620 nm to 900 nm using the F-7100 fluorescence spectrophotometer (Hitachi). A blank control of PBS was used to correct the final spectra.

### Molecule docking studies

Molecular docking simulations were conducted with the Molecular Operating Environment (MOE) software version 2014.0901 (Chemical Computing Group, Montreal, Canada). The crystal structure of Aβ40 (PDB ID: 2MVX) was downloaded from the Protein Database (www.rcsb.org). Firstly, hydrogen atoms were added to the protein and energy minimization was performed using the MMFF94 force field. Then, docking of the ligands into the active site of the Aβ was performed by the Triangle Matcher docking protocol of the MOE program. We kept the top 30 poses (based on the London dG scoring function). As the refinement method, the force field was chosen. The resulting poses were then re-scored by using the GBVI/WSA dG docking scoring function. The final top 5 poses were obtained for further evaluation. The figures were generated using Discovery Studio 2019 Client.

### In vitro bioluminescence studies

Solutions (80 μL) containing AkaLumine (10 μM), ATP (25 μM), Firefly luciferase (0.01 mg/ml) with and without Aβ aggregates (from 0.25 to 25 μM) were applied on a 96-well black plate, which was then subjected to imaging on an IVIS®Spectrum imaging system (Perkin Elmer, Hopkinton, MA), and data analysis was conducted using LivingImage® 4.2.1 software. Three duplicated wells were used for the quantification.

### In vitro bioluminescence intensity test in solutions (mimicking in vivo washing out effect)

A PBS solution (100 μL) containing AkaLumine (10 μM), ATP (25 μM), and Firefly luciferase (0.01 mg/ml), or a solution containing AkaLumine (10 μM), ATP (25 μM), Firefly luciferase (0.01 mg/ml) and Aβ aggregates (25 μM) or Aβ Oligomer (25 μM), was applied to an eppendorf tube with a 5KD filter insert. The insert was first imaged at 5 min, 10 min, 15 min, 30 min and 60 min after mixing,then the insert was centrifuged at 14000xg for 15 min and washed with 100uL PBS and centrifuged again. A solution (100 μL) containing ATP (25 μM) and Firefly luciferase (0.01 mg/ml) was added to the insert, which was subjected to imaging at 5 min, 15 min, 30 min and 60 min. The images were captured on the IVIS®Spectrum animal imaging system, and data analysis was conducted using LivingImage®4.2.1 software. Three duplicated inserts were used for the quantification.

### In vitro histological staining

A fresh brain tissue from a 24-month-old APP/PS1 mouse was fixed with 4% formaldehyde for 24 hours and transferred to a vial containing 30% sucrose PBS buffer at 4 °C until the tissue sunk. The tissue was embedded in OCT with gradual cooling over dry ice. The OCT embedded tissue block was sectioned into 25-μm slices with a cryostat. The tissue was washed with a diluting buffer (0.4% Triton X-100, 1% goat serum, 2% BSA in TBS) for 3×10 min, then blocked in 20% goat serum for 30 min at room temperature. Then the slice was incubated with primary antibody 6E10 overnight at 4°C. After washing with the diluting buffer for 3×10 min, the slice was incubated with a secondary antibody for 2 hours at room temperature. Then the slice was washed with TBS for 3×10 min. A solution of AkaLumine (25 μM) in 20% ethanol/PBS buffer was prepared as the staining solution. The brain slice was incubated with freshly prepared staining solution for 15 min at room temperature and then washed with 50% ethanol for 2×1 min, followed by washing with double-distilled water twice. Then the slice was covered with FluoroShield mounting medium (Abcam) and sealed with nail polish. Florescence images were obtained using the Nikon Eclipse 50i microscope with blue and green light excitation channels.

### In vitro mimic studies with brain tissues

A WT brain and a 5xFAD brain (12-months-old) were homogenized with 5.0 mL PBS buffer. Next, 1.0 mL of the homogenate suspension was transferred to an Eppendorf tube (n=3) and a solution of AkaLumine (40 μL, 2.5 μM) was added to the tube. The tube was vortexed and incubated for 20 min. After the incubation, half of the suspension (0.5 mL) was transferred to a new Eppendorf tube. The two tubes were centrifuged at 14000 RPM for 5 min. For one tube, the supernatant was removed, and 1.0 mL ethyl acetate was added to the pellet and vortex for 5 min. After spinning down, the ethyl acetate layer was collected and subjected to fluorescence spectra recording on a F7100 fluorometer. For another tube, the supernatant was removed. To the pellet, 1.0 mL fresh PBS buffer was added, and the resuspension was pipetted carefully until there is no clamped precipitate. After resuspending, the tube was centrifuged at 14000 RPM and 100 μL of the supernatant was collected (considered at 0 hour). The rest of the solution (include the pellet) was resuspended again and stirred for 0.5 hour (or 1-, 3-, and 5-hours). At each time-point, the suspension was spun down and 100 μL supernatant was collected. The collected supernatants were subjected to extraction with ethyl acetate (1.0 mL), and the ethyl acetate layers were subjected fluorescence spectrum recording (Ex/Em = 420/520 nm). The quantification was conducted with the fluorescence intensities at 520 nm.

### In vivo mimic mouse studies

In vivo bioluminescence imaging was performed using the IVIS®Spectrum animal imaging system. 7-month-old female balb/c mice were subcutaneously injected with Firefly luciferase (1.38 mg/ml in 50uL PBS) or mixture of Firefly luciferase (1.38 mg/ml) and Aβ40 aggregates (25uM in 50uL PBS) under the scalp. Then background images were acquired with open filter. Bioluminescence imaging was performed at 15, 30, 60, 90, 120, 180, 240 and 360 min after i.v. injection of AkaLumine (4 mg/kg, 5% DMSO and 95% PBS). Living Image 4.2.1 software (PerkinElmer) was used for the data analysis. For ocular imaging, 7-month old balb/c mice were injected with Firefly luciferase (1.38 mg/ml in 5uL PBS) or mixture of Firefly luciferase (1.38 mg/ml, 5uL PBS) and Aβ40 aggregates (25uM in 50uL PBS) into vitreous cavity. Then images were acquired with open filter, and bioluminescence imaging was performed at 5, 15, 30, and 60 min after i.v. injection of AkaLumine (4.0 mg/kg, 5% DMSO and 95% PBS).

### In vivo two-photon imaging

A 15-month-old 5xFAD female mouse was anesthetized with 2% isoflurane, and a cranial imaging window was surgically prepared as described ^63^. Before AkaLumine injection, two-photon images of capillary were acquired using 900-nm laser (Prairie Ultima) with 570 to 620 nm emission by injection of Rhodamine B isothiocyanate-Dextran (70KD). A bolus i.v. injection of AkaLumine (4.0 mg/kg in a fresh solution of 5% DMSO, and 95% PBS) was administered at time 0 min during image acquisition. The images were acquired with an emission channel of 500 to 550 nm. For imaging, we used a two-photon microscope (Olympus BX-51) equipped with a 20 × water-immersion objective (0.45 numerical aperture; Olympus) ^64^. Single Images were collected with 512 × 512-pixel resolution. For the time-course imaging, images were captured at speed of frame/2s. Total 40 minutes was recorded, including 2-minutes baseline recording (before AkaLumine injection). Image analysis was performed with ImageJ software.

### Preparation of mice with AkaLuc expression in brains

A craniotomy of 4 mm in diameter was prepared following the procedure for the above two-photon imaging. After drying the dura mater surface and ensuring that there was no bleeding, the procedure of virus injection was performed. Nanoject tips were prepared from capillary glass tubes that were pulled using a pipette puller to produce fine tips with high resistance. The sharp pipettes were filled with mineral oil and attached to a Nanoject III (Drummond Scientific, Broomall, PA). Next, most of the mineral oil was pushed out of the pipette using the Nanoject, and a vector solution was drawn into the pipette. For accurate movement of the Nanojet, a stereotaxic micromanipulator (RWD Life Science, San Diego, CA) was used. A total volume of 1.5 uL virus containing pcDNA3 Venus-AkaLuc (Titer > 1⨯10^13^ GC/mL) was injected into 5 points (300 nL/point) each mouse at 300 micro-meter depth. After completing of the injection, the surgical field was washed with saline and then the craniotomy was put back to cover the dura mater. Silicone elastomer was applied to seal the skull, and the scalp was sutured. After housing for 3 weeks, the mice were ready for bioluminescence imaging.

### In vivo Bioluminescence imaging

In vivo bioluminescence imaging was performed using the IVIS®Spectrum animal imaging system. 11-Month-old female 5xFAD/AkaLuc mice (n = 4) and age-matched female WT/AkaLuc mice (n = 3) were shaved before background imaging and were intravenously injected with a freshly prepared solution of AkaLumine (16.0 mg/kg, 5% DMSO and 95% PBS). Images were acquired with open filter or specific emission filters from 580 nm to 760 nm with an interval of 20 nm. Bioluminescence signals from the brains were recorded before and 5, 15, 30, 60, 120, 240, 360, 480 and 1440 min after i.v. injection of AkaLumine. Living Image 4.2.1 software was used for the data analysis.

### BLIAR for monitoring Aβ changes in different ages of transgenic mice

5xFAD/AkaLuc and WT/AkaLuc mice were prepared as the above (female, n=3 at the age of 8-month-old and 12-month-old, respectively) and wild-type mice (female, n=4). After 3 weeks of wound healing and virus expression, we followed the above protocol to image the 8- and 12-months-old mice. Region of interest (ROI) was drawn around the brain region and the signal at each time point was normalized by background signal. The cerebral signal was fit with two-phase decay model and the halftime (T_1/2_) of each group was quantified by using Graphpad Prism 8.

### BLIAR for monitoring therapy in transgenic mice

5xFAD/AkaLuc mice (female, n= 10) at the age of 8-month-old were prepared as the above. However, 4 mice were not survived after the surgery. After 3 weeks of wound healing and virus expression, the mice were treated with γ-secretase inhibitor Avagacetat (10mg/kg, i.p. injection) for three times a week. After 2-months treatment, the mice were imaged by following the similar imaging protocol as the above, except that i.p injection was performed (32 mg/kg, 10%DMSO, 5% cremophore, 85%PBS). After the imaging procedure, the mice were sacrificed, and brains were imaged with Venus fluorescence to measure the Aka Luciferase expression. The brains were then homogenized, and the concentration of Aβ40 and Aβ42 were measured by Meso Scale Discovery (MSD) technology with V-PLEX Plus Aβ Peptide Panel 1 (6E10) Kit according to the manufacturer protocol ^57,58 65^, and each sample was triplicated for the measurement.

## Acknowledgment

This work was supported by NIH R01AG055413, R21AG059134, and R21AG059134 awards (C.R.). Shiqian Shen acknowledges NIH R03 AG067947; R35 GM128692; R61 NS116423; and AG065606. The authors would like to thank Pamela Pantazopoulos, B.S. for proofreading this manuscript. The authors thank MGH IACUC, animal facility, and veterinary services for supporting this project. The authors thank Drs. Abbas Yaseen, Sava Sakadzic, and Caroline Magna for maintaining the two-photon microscopy facility.

## Supplemental Figures

**SI Fig.1.**
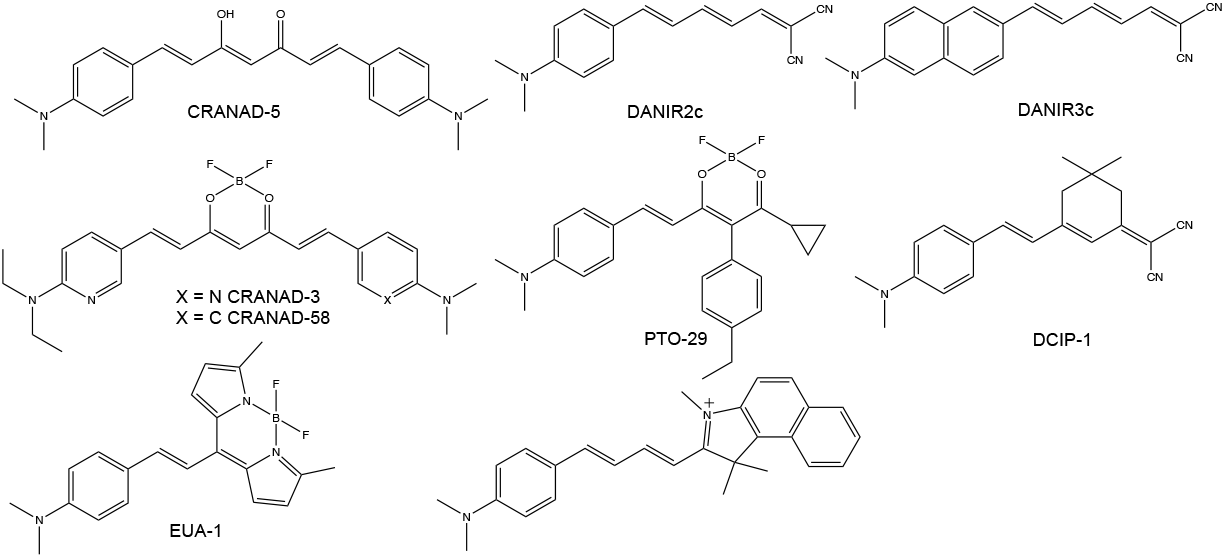
Chemical structures of Aβ fluorescence probes that have certain structural similarities to AkaLumine.

**SI Fig.2.**
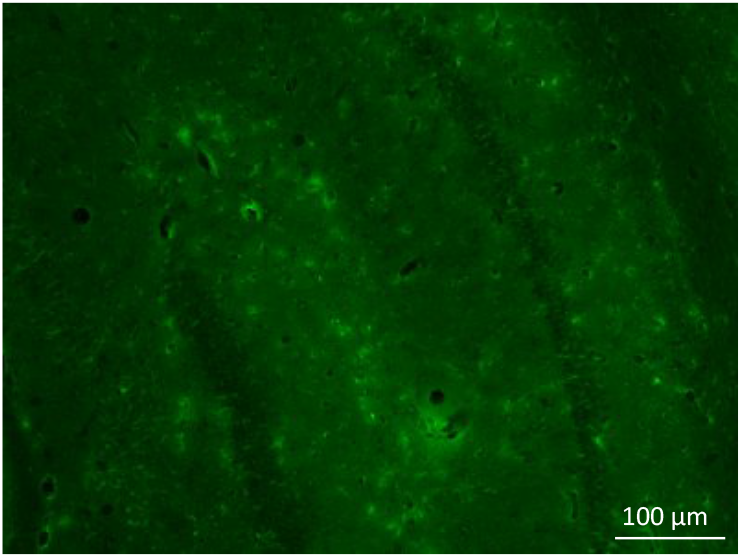
Microscopic fluorescence imaging of a brain tissue slide from a 12-month 5xFAD mouse brain with firefly luciferin. No apparent plaques could be identified.

**SI Fig.3.**
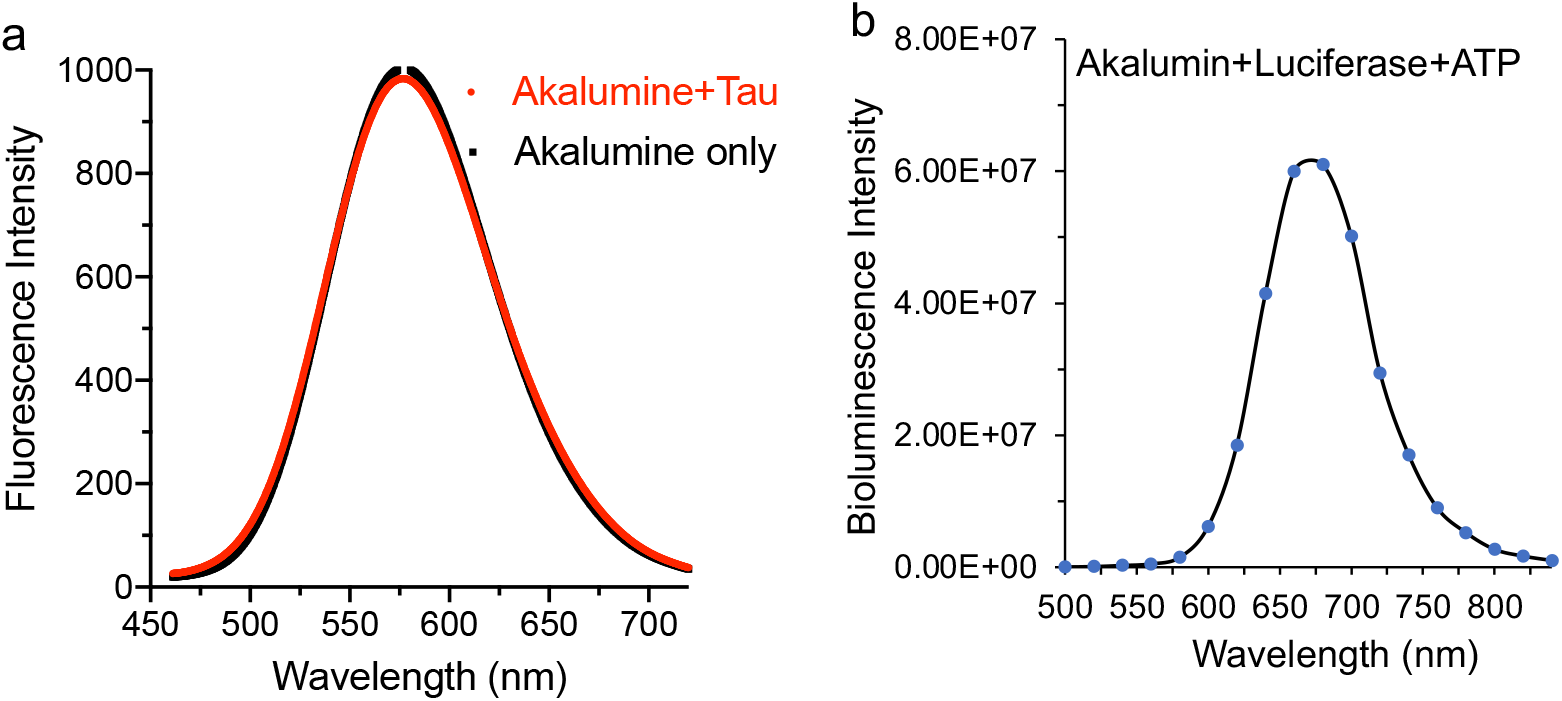
a) Fluorescence spectra of AkaLumine with and without tau aggregates. b) Emission spectrum of AkaLumine in the presence of FLuc and ATP.

**SI Fig.4.**
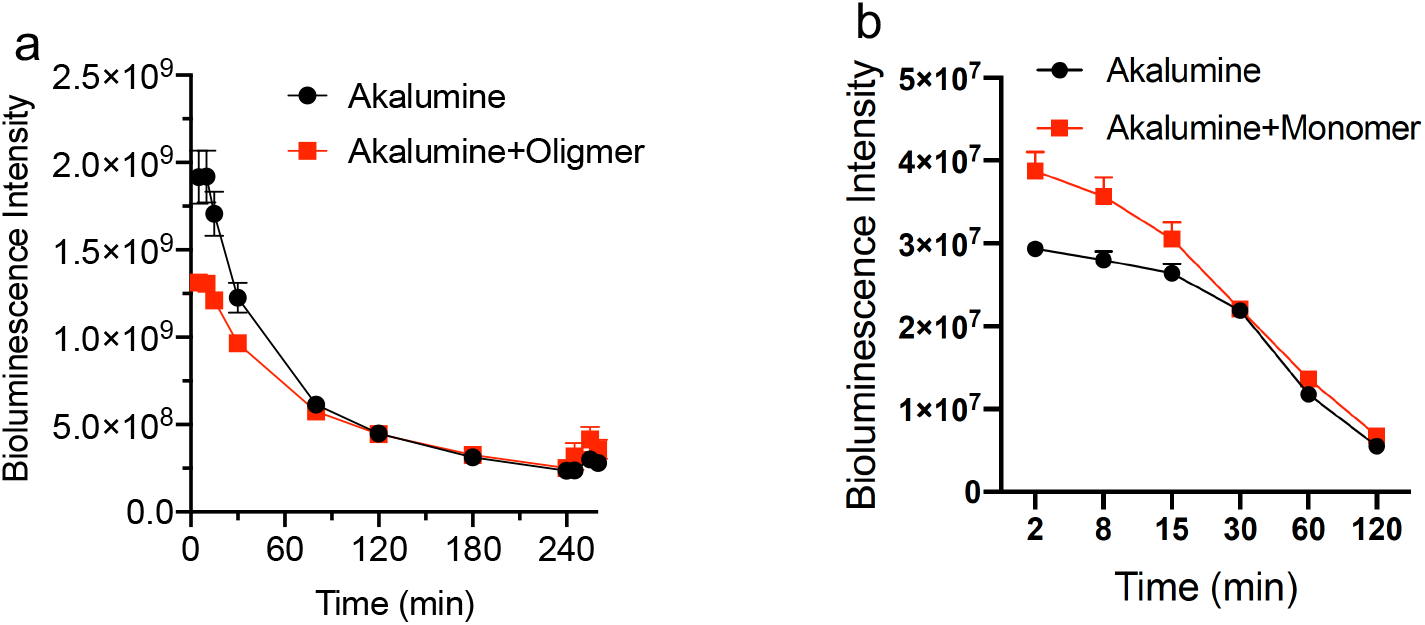
a) Full dynamics of bioluminescence intensity of oligomers with filtering (sequester + release) in the presence of AkaLumine and FLuc and ATP. b) Full dynamics of bioluminescence intensity of Aβ40 monomers in the presence of AkaLumine and FLuc and ATP.

**SI Fig.5.**
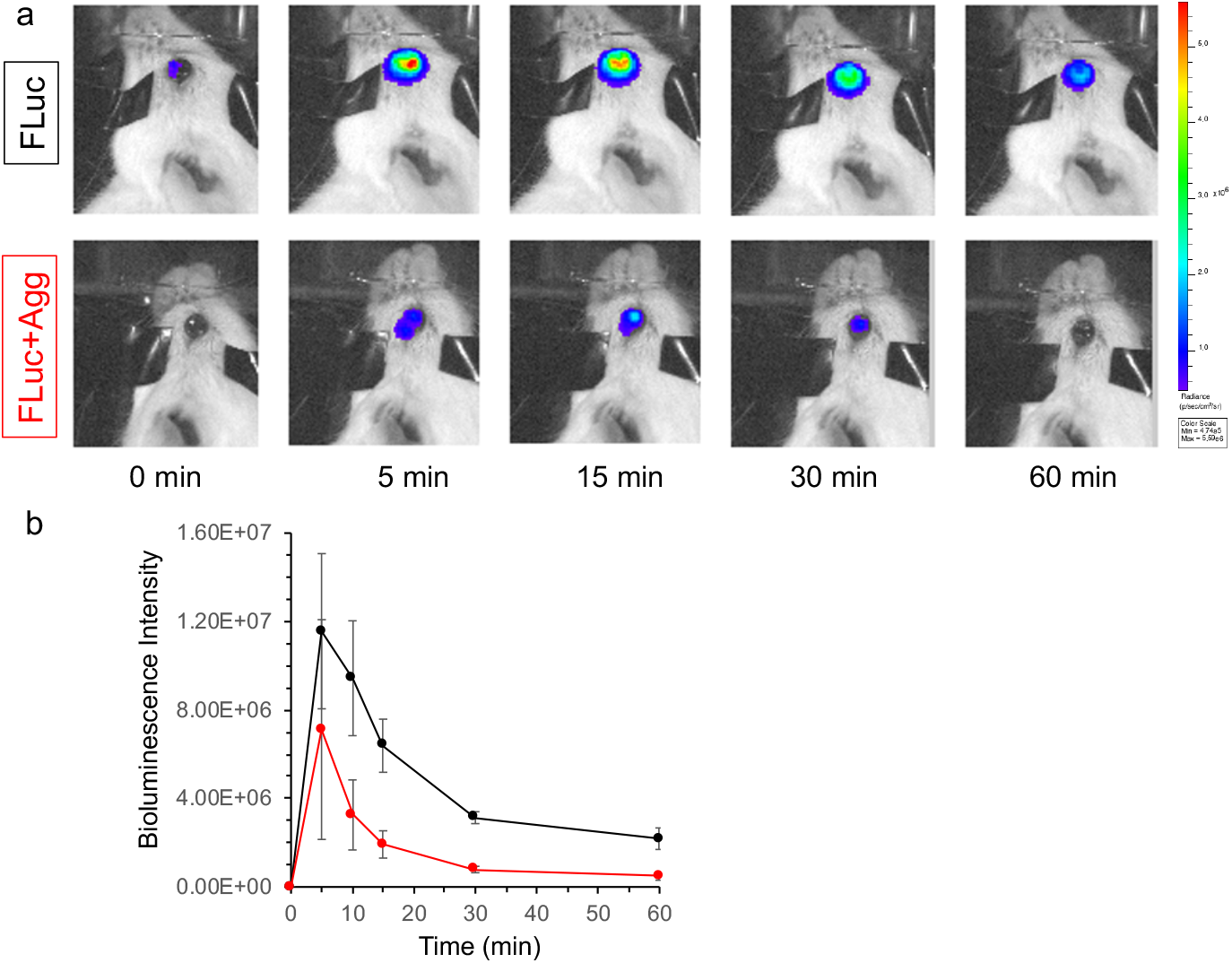
a) Representative ocular images of BLIAR after the injection of FLuc only (upper) and FLuc and Aβ40 aggregates (lower). It is clear that the presence of Aβ aggregates could lower the bioluminescence intensities. b) Quantitative analysis of BLIAR ocular images (n=3).

**SI Fig.6.**
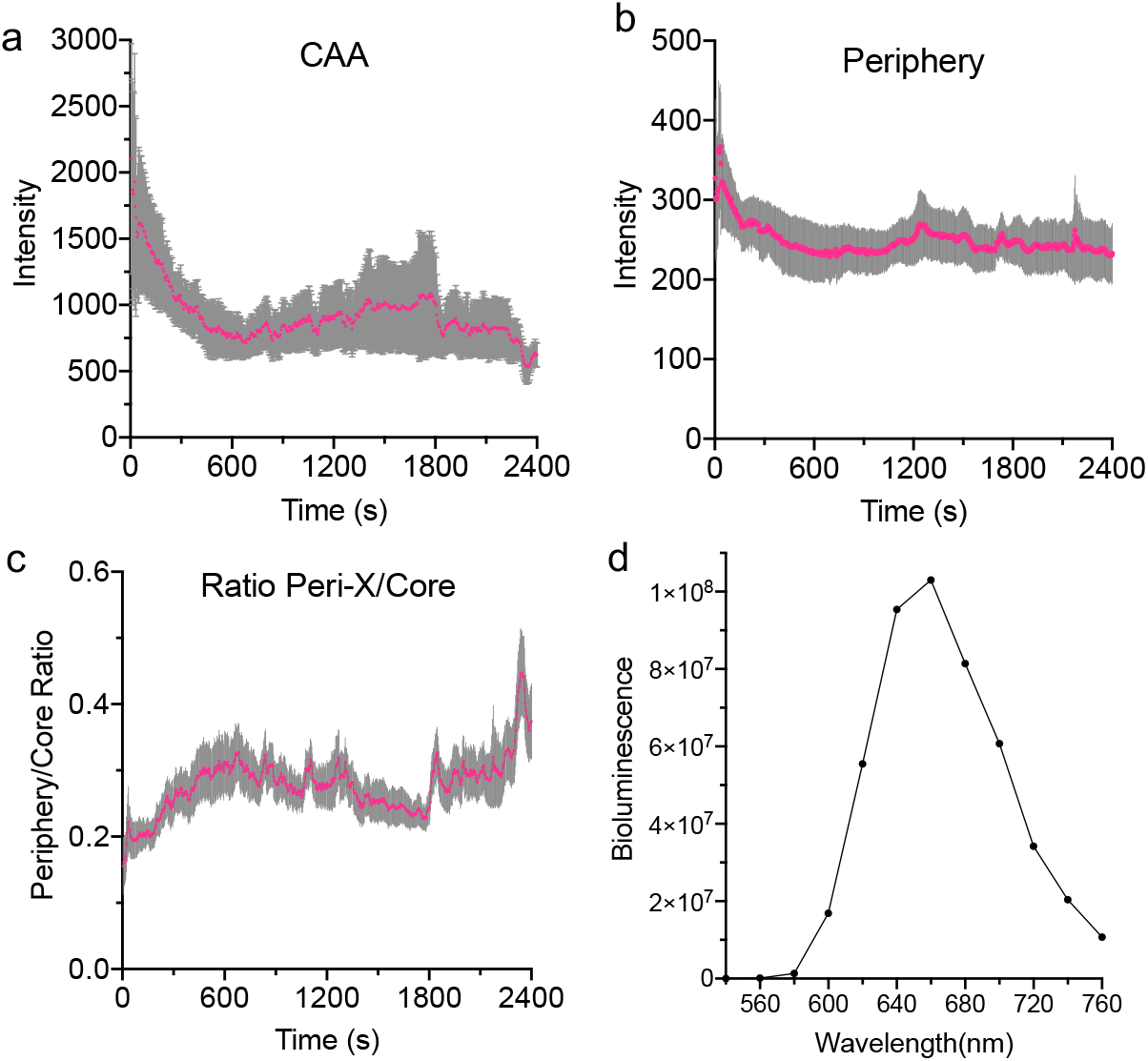
a-c) Two-photon imaging with a 5xFAD mouse. a) Time-course of CAAs (n=6). b) Time-course of peripheral areas of plaques (n=18). c) Time-course of the intensity ratio of periphery/core. d) Bioluminescence emission spectrum of AkaLumine from in vivo brain imaging.

**SI Fig.7.**
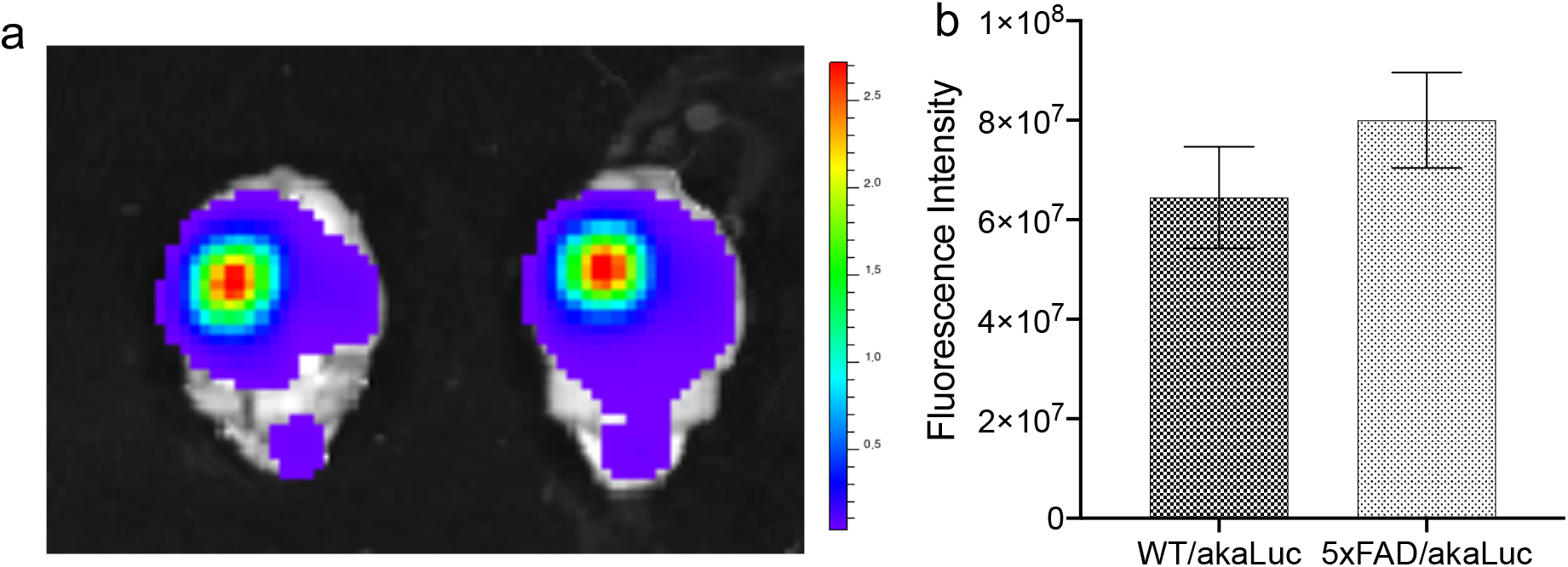
a) Representative fluorescence imaging of brains for Venus yellow proteins. b) Quantitative analysis for ex vivo brain images with the Venus gene reporter.

**SI Fig.8.**
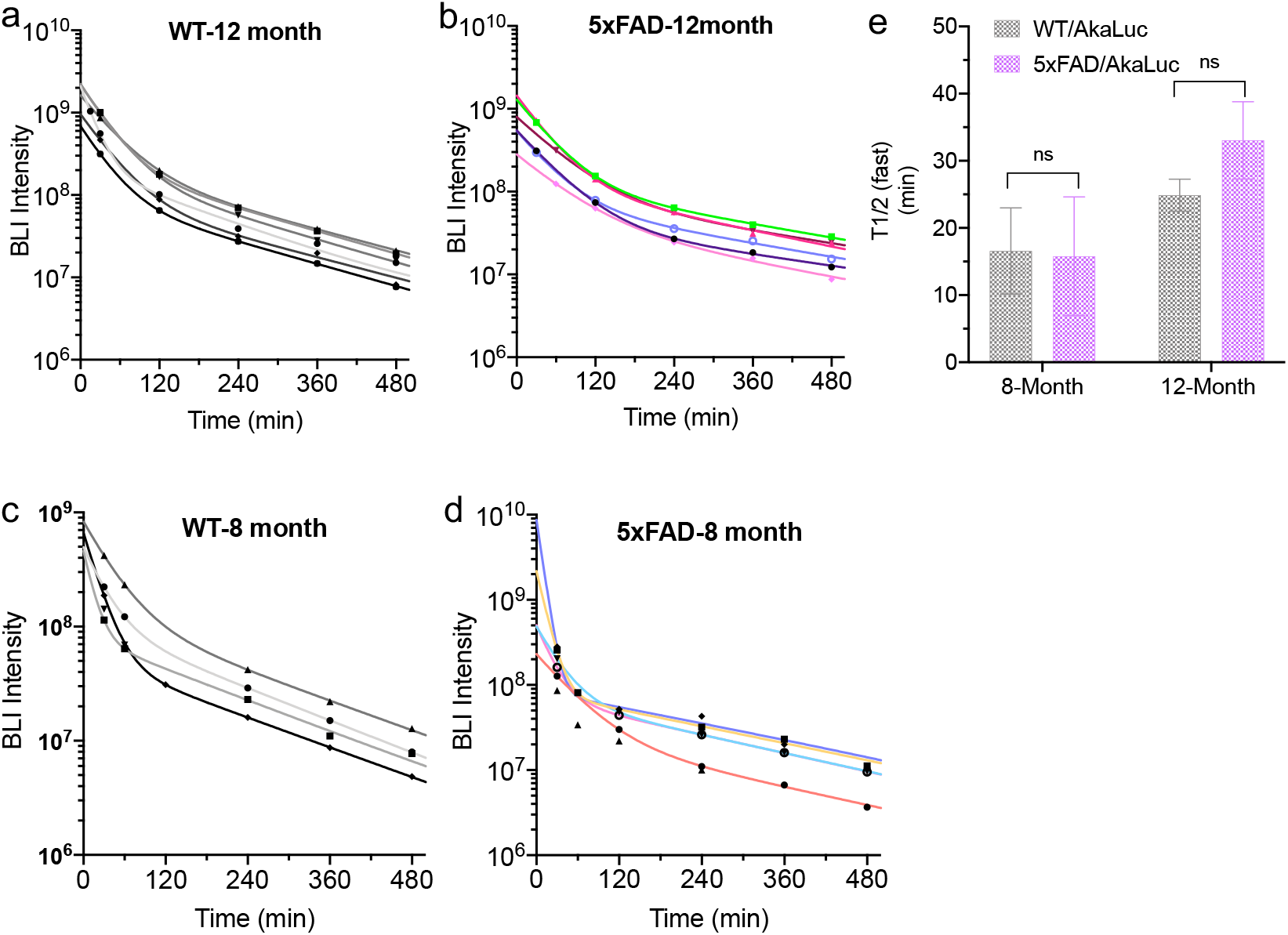
In vivo BLIAR imaging with 8- and 12-month-old WT/AkaLuc and 5xFAD/AkaLuc mice. a-d) The fitting curves with two-phase decay of the BLI intensity for 0-480 min. e) The halftime T_1/2(Fast)_of the fast decay phase from 8- and 12-month-old mice.

**SI Fig.9.**
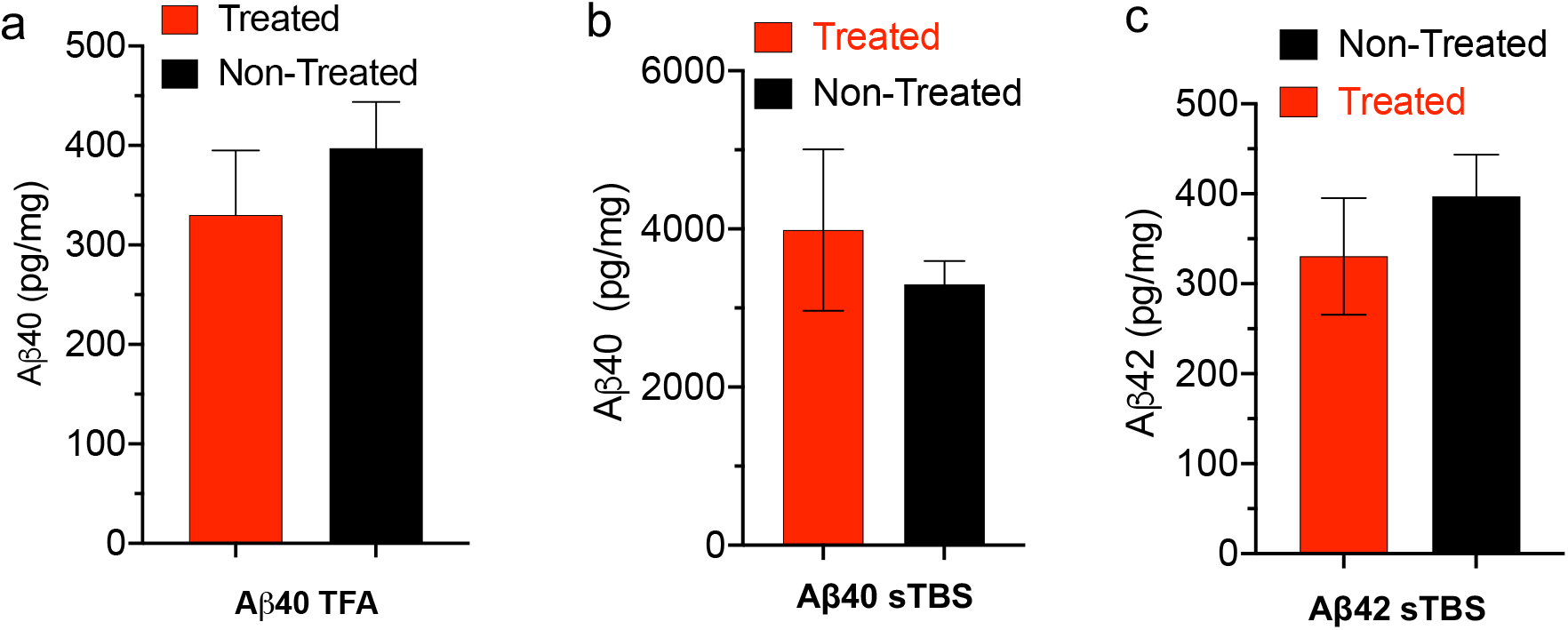
MSD measurements of Aβ40 and Aβ42 concentrations of ex vivo brain tissues after sTBS extraction and TFA extraction (n=3). a) Aβ40 concentrations in brain tissues of the treated and non-treated groups after TFA extraction. b-c) Aβ40 and Aβ42 concentrations in brain tissues of the treated and non-treated groups after sTBS extraction.

## Notes

### Competing Interest Statement

The authors have declared no competing interest.

